# Diffusion pseudotime robustly reconstructs lineage branching

**DOI:** 10.1101/041384

**Authors:** Laleh Haghverdi, Maren Büttner, F. Alexander Wolf, Florian Buettner, Fabian J. Theis

## Abstract

Single-cell gene expression profiles of differentiating cells encode their intrinsic latent temporal order. We describe an efficient way to robustly estimate this order according to a *diffusion pseudotime*, which measures transitions on all length scales between cells using diffusion-like random walks. This allows us to identify cells that undergo branching decisions or are in metastable states, and thereby genes differentially regulated at these states.

---

Cellular programs are driven by gene-regulatory interactions, which due to inherent stochasticity as well as external influences often exhibit strong heterogeneity and asynchrony in the timing of program execution. Time-resolved bulk transcriptomics averages over these effects and obscures the underlying gene dynamics. Instead, single-cell profiling techniques allow a systematic observation of a single cell’s regulatory state^1^, capturing cells at various stages in their respective process^2,3^. To infer gene dynamics and hence the sequence of cellular programs, the collective (‘universal’) process dynamics (Box 1) can be reconstructed by reordering cells according to some measure of expression similarity. This so-called pseudotemporal ordering was initially proposed for bulk expression^4^, and was later extended to single-cell RNA-seq ^5^ and protein profiles from mass cytometry^6^.

### Box 1: Universal time

In contrast to continuous time observations of a single cell e.g. from time-lapse microscopy, high-throughput snapshot experiments such as single cell RNA-seq or FACS only encode the collective (‘universal’) time dependence of cells, not the stochastic trajectories of single cells. We define universal time as the geodesic distance on the manifold that is associated with the deterministic program underlying the stochastic cellular process. For time-lapse data, universal time can be constructed by estimating the velocity *v*(*t*) tangential to this manifold *C* from local averages of single-cell trajectories. The geodesic distance

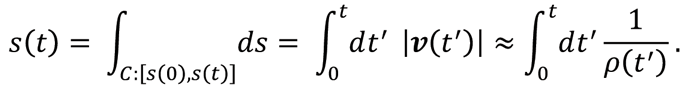

then quantifies the arc length i.e. the universal time along the manifold, where *ρ*(*t*) denotes the local density of samples on a single cell trajectory (see Supplementary Sec. 1).

Pseudotimes are proxies for universal time (Supplementary Figs. 1-3). Our proposed DPT approximates universal time better than other pseudotime schemes as it does not involve dimension reduction, and better than diffusion distance^11^ as it accounts for random walks on all length scales.

Ultimately these approaches aim to fully understand differentiation dynamics as paths on Waddington’s ‘epigenetic landscape’^7,8^. However so far it is unclear how to identify cells at critical branching decisions as well as quiescent or metastable cells, for which there is no notion of temporal ordering. Moreover, as novel experimental techniques such as droplet sequencing^9,10^ allow to profile tens of thousands of cells, there is an urgent need for computationally efficient, scalable and robust algorithms.

To overcome these problems, we introduce ‘diffusion pseudotime’ (DPT). It measures progression through branching lineages using a random-walk-based distance in diffusion map space^11^ and allows for branching and pseudotime analysis on large-scale RNA-seq data sets. Even in the absence of branching, DPT is significantly more robust with respect to noise in low-density regions and cell outliers than existing methods, which rely on the estimation of minimum spanning trees^5^ or sampling-based distances^6,12^.

Diffusion pseudotime is computed in three steps (Fig. 1a and Online Methods). First, a transition matrix ***T*** that approximates the dynamic transitions of cells through stages of the differentiation process is determined. The right eigenvectors of ***T*** are known as diffusion components and have been used in diffusion maps for visualizing single cell RNA-seq data^13,14^. While using only few diffusion components yields interpretable visualizations, important information may be lost by removing the others. Consequently, DPT is based on the full rank **T** rather than a low rank approximation.

**Figure 1:**
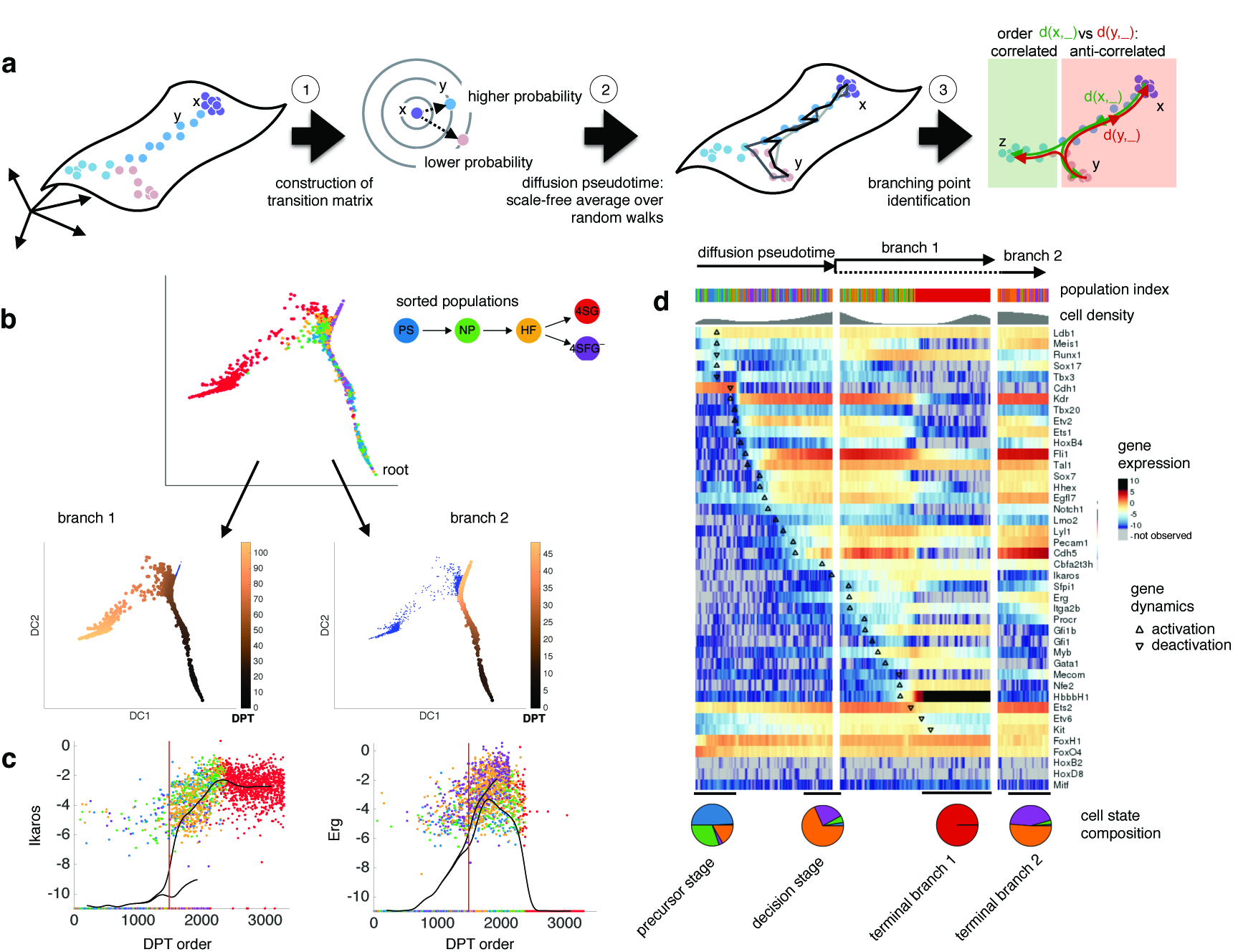
Diffusion pseudotime reveals temporal ordering and cellular decisions on the single cell level. **(a)** Outline of the computational workflow. The diffusion transition matrix ***T***_*xy*_ is constructed by superimposing local kernels at the expression levels of cells *x* and y (1). The diffusion pseudotime dpt*(x,y)* approximates the geodesic distance of *x* and *y* on the mapped manifold (2). Branching points are identified as points where anti-correlated distances from branch ends become correlated (3). **(b)** Application of DPT to single-cell qPCR of 42 genes in 3934 single cells during early hematopoiesis^15^, sorted from 5 different populations: primitive streak (PS), neural plate (NP), head fold (HF), four somite GFP negative (4SG-), four somite GFP positive (4SG+). DPT identifies one branching point. **(c)** Exemplary dynamics of genes Erg and Ikaros show qualitatively different behavior in the two branches, black lines describe a moving average over 50 adjacent cells along the respective branch. **(d)** Heatmap of gene expression, with cells ordered by diffusion pseudotime and genes ordered according to the onset of first major change in expression (see Supplementary Sec. 7.2, smoothed along 50 adjacent cells, see Supplementary Fig. 5 for non-smoothed version). Bars on top indicate the cells’ population (b) and cell density, respectively, with high density regions indicating metastable states. The time series were clustered temporally by the time point of the first transition event (precursor branch, branch 1 and none, respectively, Supplementary Fig. 6). The pie charts (bottom) show the fraction of cells making up the metastable states (black horizontal line).

In the second step, the distance dpt(*x,y*) between two cells with index *x* and *y* is computed as

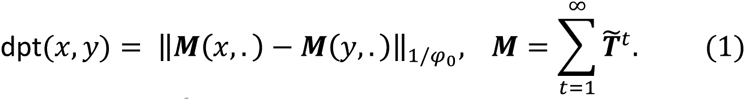

Here, ‖… ‖_1/*φ*_0__ denotes the *L*^*2*^ norm weighted by the first left eigenvector *φ*_0_ of ***T***, the steady state (Online Methods, Supplementary Sec. 1.3). Instead of the probability (***T***^*t*^)_*xy*_ for a random walk of fixed length^11^ *t* from x to y, in Eq. (1), we compute the accumulated transition probability (***M***)_*xy*_ of visiting y when starting from x over random walks of all lengths. This is done using the modified transition matrix 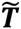, which is defined as ***T*** without the eigenspace associated with the steady state *φ*_0_, and therefore describes how the steady state is approached. Fixing a known root cell *r* as start of the dynamical process of interest, we define the pseudotime of cell *x* as dpt*(x,r)*.

In the third step, branching points are identified by comparing two random walks over cells, one starting at the root cell *r* and the other at its maximally distant cell *y*, measuring pseudotimes with respect to *r* and *y*, respectively. The two sequences of pseudotimes are anticorrelated until the two walks merge in a new branch, where they become correlated (Online methods). This criterion robustly identifies branching points as we illustrate for simulation data for which the ground truth is known (Supplementary Fig. 4).

To illustrate the ability of DPT to identify branches on real data, we reanalyzed a single-cell qPCR data set focusing on early blood development^15^, for which we have shown previously that diffusion maps allow to visually identify a precursor branch splitting up into two separate lineages (cf. Fig. 1b). DPT ordered cells along their developmental trajectory and identified cells at the branching point. The three identified branches qualitatively coincide with a precursor branch and the reported blood (branch 1) and endothelial branches (branch 2)^15^. In particular we identified characteristic patterns in the developmental stages in blood progenitors (Fig. 1d), namely the hemangioblast-like sequence^16^ (subsequent up-regulation of *Cdh1* to *Tal1* and *Cdh5)* at the precursor branch^15^ and the endothelial differentiation route^15^ on branch 2 (elevated levels of *Pecam1, Erg* and *Sox17* amongst others). Further, we find the erythroid-like sequence of *Etv2, Tal1, Runx1* and *Gata1^17^* at branch 1. The temporal resolution obtained by DPT indicates immediate (directly after branching, cf. *Ikaros* expression in Fig. 1c) and late transitions (cf. *Erg* in Fig. 1c) as well as a number of intermediate regulatory events^15^ until the onset of *Hbb-bH1* expression (cf. Fig. 1d, black triangles), hinting at potential novel regulatory interactions.

DPT identified regions of small time-steps i.e. of high cell density (Fig. 1d, top and Supplementary Fig. 7b) along the differentiation process. These high-density regions indicate metastable states, which correspond to biologically meaningful intermediates: We found four metastable states with expression patterns of precursor cells, hemangioblast-like cells at the decision state, erythroid-like and endothelial-like cells. Notably, both decision and precursor states consist of cell mixtures from two or three different stages, stressing the asynchrony of developmental stages that could not be resolved without pseudotemporal ordering.

To identify key decision genes, we quantified expression differences in the identified states of decision versus precursor using MAST^18^ (Supplementary Fig. 8a). This resulted in more than 50% of changed genes (27 out of 42), including *Pecam1* and *Cbfa2t3h*, which are known to indicate hematopoietic and endothelial development^16^, respectively. In contrast, only 24 genes are differentially expressed between sorted cells from head fold and primitive streak, all changing gradually but preserving bimodal distribution (Supplementary Fig. 8d and Supplementary Table 1). Also, differential gene expression between HF and 4SG-cells fails to identify endothelial differentiation but brings up erythroid factors (*Runx1, Ikaros* and *Gfi1b* amongst others, see Supplementary Fig. 8e and Supplementary Table 2). In summary when comparing differentially expressed genes between metastable states, we identify more genes than comparing developmental stages, and the genes have less bimodal expression. This shows that the anatomical stages overstate developmental heterogeneity thus disguising the role of key factors.

DPT copes well with large-scale experiments such as scRNA-seq combined with droplet barcoding^10^: In the experiment, Klein et al. monitored the transcriptomic profiles and heterogeneity in differentiation of mouse ES cells after LIF withdrawal (Fig. 2a). After cell-cycle normalization (Supplementary Figs. 9-10), DPT describes a single differentiation path from which two populations branch off. With increasing pseudotime, we observe upregulated epiblast markers (*Krt8/18/19*) and downregulated pluripotency factors (*Nanog*, Fig. 2b). Clustering of the gene expression dynamics identified four clusters with different temporal behaviors (Fig. 2c and Supplementary Fig. 11) but coherent biological functions (Fig. 2d). Early pseudotime coincides with day 0 cells exhibiting strong expression of pluripotency factors (purple cluster). Then, a small subpopulation (57 cells) mainly consisting of day 2 cells branches off, enriched in neuron-associated genes *(Bc1, Lin7b, Snord64, Tagln3, Dtnbp1*, Nenf; 6 out of 22, see Supplementary Fig. 12). Subsequent stages are characterized by a gradual decrease of pluripotency factors and slow rise of both primitive endoderm markers (yellow cluster) and epiblast markers (orange cluster)^19^. In late pseudotime, another two branching event gives rise to a population with increased primitive endoderm markers (21 cells), whereas epiblast marker genes rise two-to three-fold in the other branch (120 cells). Altogether, DPT is able to remove asynchronity of scRNA-seq snapshot data from several days, aligning cells in terms of their degree of differentiation.

**Figure 2:**
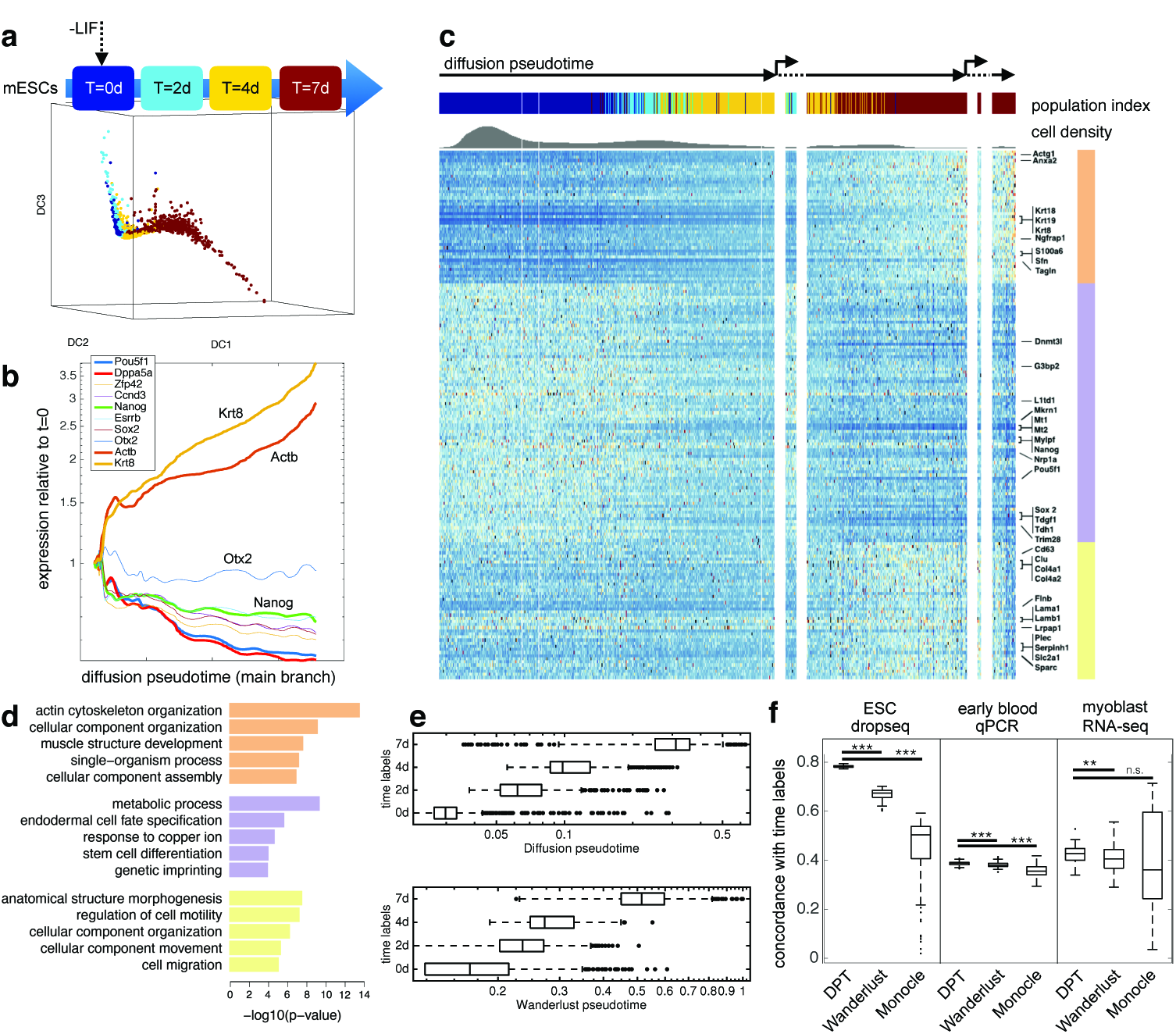
Diffusion pseudotime identifies differentiation dynamics in droplet-based scRNA-seq experiments^10^. **(a)** Mouse ESCs after LIF withdrawal were harvested at T=0, 2, 4 and 7 days and profiled with the dropSeq protocol, giving 2717 cells with 24175 observed unique transcripts^10^. After correction for cell cycle variation, a low dimensional visualization using diffusion maps shows overlapping but directed temporal dynamics. **(b)** Temporal dynamics of selected genes as in the original publication reconstructed by DPT, relative to expression at initial time point. **(c)** DPT identifies a main time axis with two minor branching events: an early side branch and late separation of cells enriched with markers for epiblasts and primitive endoderm (top). Population and cell density are shown as in Fig. 1d. The heatmap depicts gene dynamics after hierarchical clustering and removal of a fluctuating, mainly constant subgroup (cf. Supplementary Fig. 11b): The dynamic subgroups (indicated by color bar, right) consist of epiblast markers such as Krt8/18/19, Sfn, Tagln (orange), gradual downregulated pluripotency factors such as *Pou5f1 (Oct4), Sox2, Trim28, Nanog* (purple) and slow consistent upregulated primitive endoderm markers such as *Col4a1/2, Lama1/b1, Serpinh1, Sparc* (yellow). **(d)** Gene ontology enrichment shows a cellular reorganization signature (orange), a metabolic signature consistent for differentiation (purple) and a cell motility signature (yellow). **(e)** Pseudotime distribution of cells in the experiments from the four different days, for DPT and Wanderlust. Diffusion pseudotime orders cells well along the four temporal categories (Kendall rank correlation 0.78±10^−3^), significantly better than pseudotemporal ordering by Wanderlust (Kendal rank correlation 0.71±10^−3^, see also Supplementary Fig. 13). **(f)** Comparisons of Kendall rank correlation on bootstrap samples (n=100 bootstrap runs, downsampling to maximal 1800 cells for Wanderlust and DPT, 700 cells for Monocle due to performance issues) for the presented ESC data set, the qPCR data set from figure 1 and an scRNA-seq of differentiating myoblasts from the Monocle paper^5^ show that DPT consistently outperforms the other two methods (2-sided t-test with significance levels *p<0.05; **p<0.01; ***p<0.001, n.s. not significant).

To evaluate DPT’s performance without branching compared to Wanderlust and Monocle, we counted how often a pseudotime puts a cell from a later temporal sorting before an earlier one (measured by Kendall rank correlation *τ*). DPT reconstructs the temporal orders of ESC differentiation with significantly higher accuracy than Wanderlust (*τ* = 0.78±10^−3^ versus 0.71±10^−3^, respectively, Fig. 2e and Supplementary Fig. 13). This holds true also when compared to Monocle in repeated bootstrap runs and on other data sets (Fig. 2f and Supplementary Table 3).

In conclusion, we introduce DPT as a pseudotime measure that overcomes the deficiencies of existing approaches: it is able to deal with branching lineages and identifies metastable or steady states, it is statistically robust, and its computation can be scaled to large datasets without dimension reduction. Compared to Wanderlust^6^, which has been proposed for the lower-dimensional mass cytometry data, we replaced approximate and computationally costly sampling of shortest paths by the exact and computationally cheap average over random walks in eq. (1). Compared to Monocle^5^, which works on RNA-seq data but only after dimension reduction and on medium sample numbers, DPT’s average over all random walks is significantly more robust than Monocle’s minimum spanning tree approach (Fig. 2e).

In the future, robust computation of pseudotimes will allow inferring regulatory relationships with much higher confidence than based on perturbations alone^15^, and we expect DPT to allow scaling this to genome-wide models. Recently pseudotemporal ordering has been applied to cell morphology to identify cell cycle states^20^ – here diffusion pseudotime would allow inclusion of branching for example to identify cells switching into a quiescent state as well as comparison to time-lapse microscopy via universal time. To summarize, diffusion pseudotime provides a powerful and robust tool to order cells according to their state on differentiation trajectories in single-cell transcriptomics studies.

## Acknowledgements

We would like to acknowledge Carsten Marr, Jan Hasenauer, Matthias Heinig, Jan Krumsiek and Thomas Blasi for their helpful advice and comments on the manuscript.

M.B. is supported by a DFG Fellowship through the Graduate School of Quantitative Biosciences Munich (QBM). F.A.W. acknowledges support by the “Helmholtz Postdoc Programme”, Initiative and Networking Fund of the Helmholtz Association. F.B. is supported by the UK Medical Research Council (MRC) via a Career Development Award (Biostatistics). F.J.T. acknowledges financial support by the German Science Foundation (SFB 1243 and Graduate School QBM) as well as by the Bavarian government (BioSysNet).

## Methods

Methods and any associated references are available in the online version of the paper.

## Accession codes

A MATLAB implementation of DPT is available on http://www.helmholtz-muenchen.de/icb/dpt.

## Author contributions

L.H. developed the method and the computational tools, performed the analysis and wrote the paper. M.B. contributed to the analysis of results. F.B. and F.A.W. helped interpret the results and write the supplement. F.J.T. conceived and supervised the study, contributed to the method development and wrote the paper with help from all co-authors.

## Competing financial interests

The authors declare no competing financial interests.

## Online Methods

### Overview of DPT algorithm

0) (Initialization) inputs the following:

a. The n by G data matrix
b. One (or several) root cell(s).
c. Diffusion maps options “classic” or “locally scaled” and respectively the parameters “θ” (kernel width) or “κ” (numberof nearest neighbours for adjusting the kernel width).
  1. Computes the transition matrix *T.*
  2. Builds the accumulated transition matrix M and computes diffusion pseudotime with respect to the specified root. If several roots are defined, DPT averages the pseudotime for each cell y over these roots.
  3. DPT iteratively assigns cells to branches and subbranches. DPT groups the cells for each branch and provides diffusion pseudotime for each group.

### Diffusion pseudotime

We calculate the diffusion maps transition matrix ***T*** and its right and left eigen-vectors Ψ_0_ and *φ*_0_. It then computes the accumulated transition probabilities over all numbers of time steps.

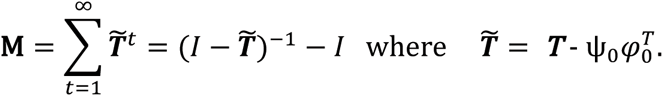

This is done relative to the steady state *φ*_0_, which stores no information about dynamics. Fixing a known root cell *x* as start of the dynamical process of interest, Diffusion pseudotime of cell y is defined as a density weighted L^2^ norm

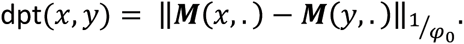

Further details are given in Supplementary Sec. 3.

### Branch assignment

We find the cell *y* with the maximal dpt distance from the root(s) *x* and also another cell *z* which has maximal distance to *x* and *y*:

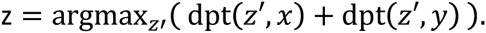

If the manifold is branching, then as defined *y* and *z* will provide cells at two different tips of two branches.

DPT then obtains two orderings *O*_*y*_=dpt(.,*y*) and *O*_*z*_=dpt(.,*z*) and determines the cutoff cell until which the sequence of ordered cells in *O*_*x*_ (call them *X*_*i*_), *O*_*y*_ and *O*_*z*_ become maximally correlated using Kendall’s rank correlation. DPT thus assigns cells *X*_*i*_ to the branch of *x*.

DPT treats *y* and *z* as root of the subbranches *Y*_*i*_ and *Z*_*i*_ respectively and in a similar way searches for new subbranches within each branch. Further details are provided in Supplementary Sec. 4.

### Metastable states

The pseudotemporal ordering of cell populations reflects gradual and switch-like changes along a certain branch. Highly similar cells have small distance in the gene space and a high probability to be reached by a random walk as defined by the transition matrix *T*. Then, the difference in pseudotime between such cells is small, i.e., the density of the distance to the root cell *dpt(r,.)* increases at sites, where highly similar cells are found. In particular, developmental steady states have high densities in the pseudotime measure, but these accumulation sites are not sufficient to depict a steady state. However, these accumulation sites are not sufficient to depict a steady state.

### Detecting transcriptional changes

To identify the succession of switch-like transcriptional changes revealed by the pseudotemporal order in qPCR data, we computed an approximate derivative of the smoothed gene expression level along branch 1. A switch-like change is defined as the maximum in the derivative (details in Supplementary Sec. 7.2).

### Differential expression analysis

We employed a two-part, generalized linear model that allows to quantify the proportion of cells expressing a certain gene as well as the mean expression level, a modified Hurdle model^1^. Briefly, the model has two parts: A discrete part to decide whether a gene is expressed and a continuous part using a normal distribution if the gene is expressed. Then, a likelihood ratio test is used to identify differentially expressed genes (details in Supplementary Secs. 5 and 7.3. and Finak et al^1^).

### ESC qPCR Data

We reanalyzed a single-cell qPCR dataset (normalized version with 3934 cells, 42 genes) focusing on early blood development^2^. For each gene, the limit of detection (LOD) was the average Ct value for the last dilution at which all six replicates had positive amplification. The overall LOD of 25 for the gene set was the median Ct value across all genes. Raw Ct values and normalized data can be found in Supplementary Table 7 of Moignard et al^2^. Gene expression was subtracted from the limit of detection and normalized on a cell-wise basis to the mean expression of the four housekeeping genes (*Eif2b1, Mrpl19, Polr2a* and *Ubc)* in each cell. Cells that did not express all four housekeeping genes were excluded from subsequent analysis, as were cells for which the mean of the four housekeepers was ±3 s.d. from the mean of all cells. A dCt value of −14 was then assigned where a gene was not detected. 85-90% of sorted cells were retained for further analysis. *Gata2* did not amplify correctly and *HoxB3* was not expressed in any cells, so these factors have been excluded from the analysis. The analyses were done on the dCt values for all transcription factors and marker genes, but not housekeeping genes.

### DropSeq data

We reanalyzed a single-cell RNA-seq data set using the dropSeq protocol from Klein et al^3^. Here, single cells along with a set of uniquely barcoded primers were capture in tiny droplets and sequenced. The capabilities of this technique were demonstrated using an undirected differentiation process of mouse embryonic stem cells upon leukemia inhibitory factor (LIF) withdrawal. The data set is publicly available under the GEO accession number GSE65525. Count data were normalized by library size and log_10_ transformed (see Supplementary Sec. 8.1). We corrected for cell-cycle and batch effects using scLVM^4^ on 2044 highly variable genes (see Supplementary Table 3 in Klein et al^3^). Then, diffusion map with local density rescaling (Supplementary Sec. 2) visualizes the temporal order for all cells. Hierarchical clustering was performed in R (http://www.r-project.org/) using the *hclust* package on quantile-normalized data (Supplementary Sec. 8.2) and displayed with *ComplexHeatmap* package, where the distance was defined as 1 – correlation between all samples (Supplementary Sec. 8.3). In addition, we performed a rank sums test on the first side branch to identify genes being uniquely different from initial pluripotent and late epiblast-like cells (Supplementary Sec. 8.4).

### Concordance of pseudotime with time labels

We subsampled sets of ˜70% of data and for each set performed Wanderlust^5^, Monocle^6^ and DPT pseudotime orderings. Since Monocle does not perform on very large number of cells (=10^3^), we reduced the subsampling to 700 cells when necessary. The concordance for each subset was then measured as Kendall tau correlation of each pseudotime with time labels of that subset. We then performed a t-test and calculated p-values between the histogram of the concordance measure for Wanderlust and Monocle compared to the DPT pseudotimes. The result is shown in Fig. 2f of the main text and Supplementary Table 3.

